# A Diaphanous and Enabled dependent asymmetric actin cable array repositions nuclei during Drosophila oogenesis

**DOI:** 10.1101/2020.09.29.319533

**Authors:** Gregory Logan, Brooke M. McCartney

**Affiliations:** Department of Biological Sciences, Carnegie Mellon University, 4400 Fifth Ave, Pittsburgh, PA 15213

**Keywords:** Drosophila, Oogenesis, Actin Cables, Nuclear Positioning, Enabled, Diaphanous

## Abstract

Cells reposition their nuclei for a diversity of specialized functions through a wide variety of cytoskeletal mechanisms. To complete oogenesis, Drosophila nurse cells employ novel actin cable arrays to reposition their nuclei. During oogenesis, 15 nurse cells connected by ring canals contract to “dump” their cytoplasmic contents into the oocyte. Just prior to dumping, actin cables initiate from the nurse cell cortex and elongate toward their nuclei, pushing them away from the ring canals to prevent obstruction. How the actin cable arrays generate directional nuclear movement is not known. We found regional differences in the actin cable growth rate that are dependent on the differential localization of the actin assembly factors Enabled (Ena) and Diaphanous (Dia). Mislocalization of Ena resulted in actin cable arrays with a uniform growth rate. In the absence of growth rate asymmetry, nuclear relocation was significantly altered and cytoplasmic dumping was incomplete. This novel mechanism for nuclear repositioning relies on the regulated cortical localization of Dia and Ena producing asymmetric actin cable arrays that push the nuclei away from the ring canals, enabling successful oogenesis.

**Summary statement:** This work demonstrates that an asymmetric actin cable array regulated by the differential localization of Diaphanous and Enabled is necessary to reposition nurse cell nuclei and complete oogenesis in Drosophila.

## Introduction

Regulated nuclear movements occur in a wide variety of cells, and diverse cytoskeletal-based mechanisms underlie these movements. Many contexts employ microtubule pushing (via microtubule growth) or pulling (via dynein) forces, including pre-mitotic nuclear movement in yeast (Gundersen and Worman, 2013), nuclear movement during muscle development (Azevedo and Baylies, 2020), and nuclear repositioning in the Drosophila oocyte (Tissot et al., 2017). The actin cytoskeleton is also repositions nuclei via diverse mechanisms including actomyosin contraction during neuronal migration (Nakazawa and Kengaku, 2020), and Myosin-based coupling to actin cables that relocates nuclei in migrating cells (Zhu et al., 2018). The nurse cells that promote oogenesis in Drosophila use a unique mode of nuclear repositioning: enormous arrays of actin cables that push and thereby directionally relocate nuclei in the final stages of oogenesis (Cant et al., 1994; Guild et al., 1997; Huelsmann et al., 2013; Mahajan-Miklos and Cooley, 1994). While this actin cable-driven nuclear movement has been described (Huelsmann et al., 2013), it is not known how the cable array directionally relocates the nurse cell nuclei.

The developing Drosophila oocyte is ‘nursed’ by 15 nurse cells connected to each other and to the oocyte by actin-based tunnels called ring canals (Fig 1A). The nurse cell-oocyte cluster is surrounded by a layer of somatic squamous and columnar epithelial follicle cells (Fig 1A). Through the ring canals, nurse cells send mRNAs, proteins, and organelles into the oocyte, increasing its volume ~90,000x (King, 1971). During a late stage of oogenesis (stage 11), nurse cell cortical contraction rapidly expels their remaining cytoplasmic contents through the ring canals and into the oocyte, roughly doubling the oocyte size (Fig 1A”). During stage 10B, just prior to this cytoplasmic “dumping”, striking arrays of cytoplasmic actin cables begin polymerizing at the nurse cell cortex, growing inward to contact the nurse cell nuclei (Guild et al., 1997, Fig. 1A’). These actin cables are highly ordered bundles of bundles of filaments: small 2-4 μm unit bundles containing approximately 26 parallel filaments are bundled together in an overlapping pattern to form mature cables that may extend 30 μm or more (Guild et al., 1997). Without the actin cable arrays, as in mutants of the actin bundlers Singed/Fascin (Cant et al., 1994) and Quail/Villin (Mahajan-Miklos and Cooley, 1994), the nuclei clog the ring canals during dumping and prevent the completion of oogenesis. Live imaging of this process revealed that the actin cables push the nuclei away from the ring canals as dumping commences (Huelsmann et al., 2013). In addition to the exquisite stage-specific temporal control of cable assembly, the directional repositioning of the nuclei away from ring canals also suggests spatial control of cable growth; if cables initiated simultaneously and grew at an equivalent rate from all surfaces of the nurse cell cortex, the nuclei might remain static, rather than being moved. This model predicts that the actin cable array has additional regulated temporal and/or spatial asymmetry to promote robust directional nuclear relocation.

**Figure 1:**
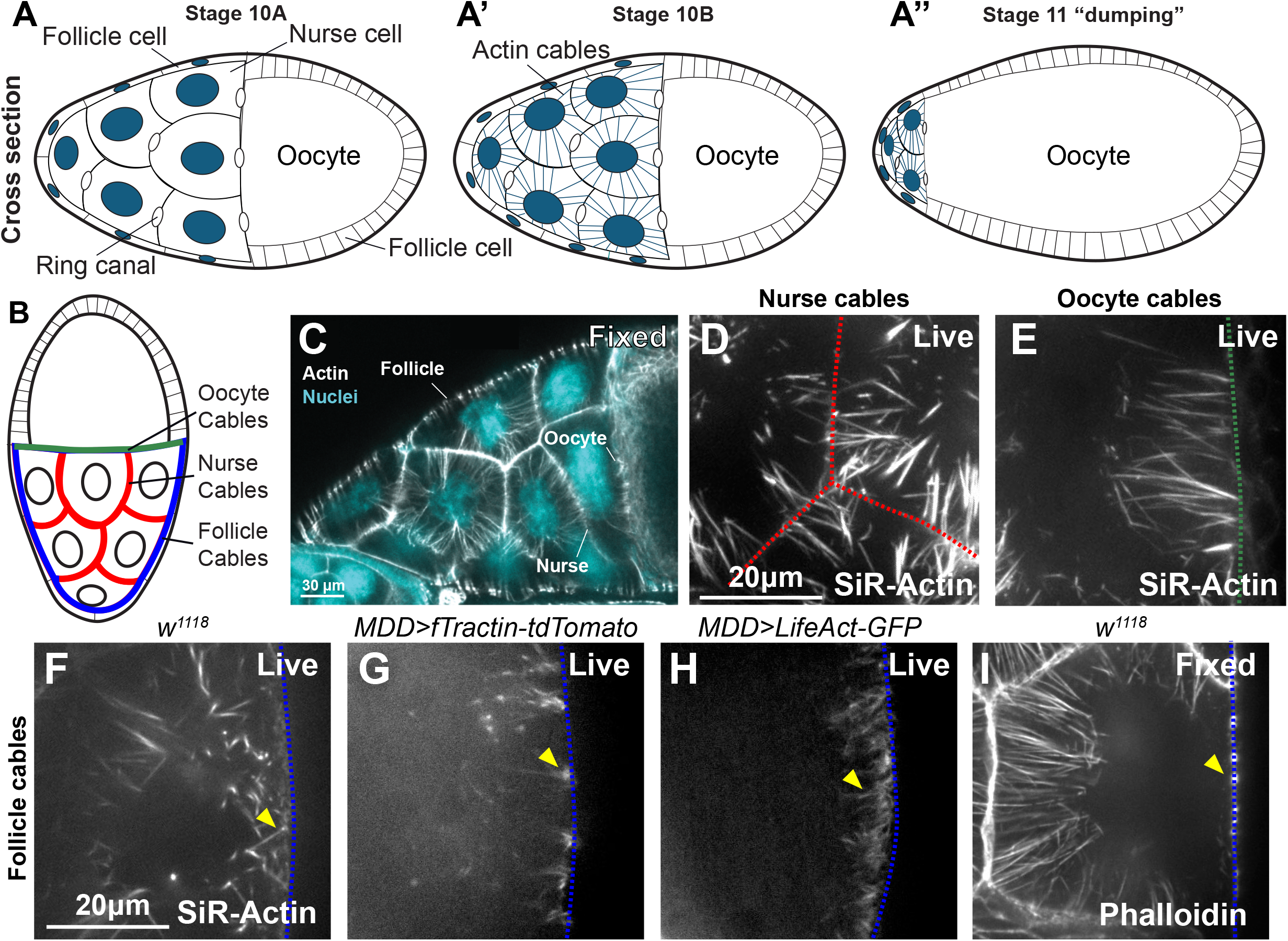
Stage 10B Drosophila nurse cells extend actin cables from all regions of the cortex. (A-A”) Schematic of stage 10-11 egg chambers showing nurse cells connected to the oocyte through ring canals. (B) Schematic of the location of the three nurse cell cable types, designated by the type of cell that the nurse cell borders. (C) A fixed egg chamber stained with DAPI and phalloidin showing the three nurse cell cable types. (D-I) Colored lines correspond to those in B, and indicate the type of nurse cell border. (D,E) Live images of SiR-Actin labeled nurse cables (D), and oocyte cables (E). (F-H) Live images of follicle cables labelled with SiR-Actin (F), fTractin-tdTomato (G), and LifeAct-GFP (H). (I) Image of a fixed egg chamber preserving no follicle cables.

Nurse cells share ring canals with adjacent nurse cells and with the oocyte, but not with the follicular epithelium (Fig 1A). The actin cables are present exclusively in the nurse cells, and for simplicity we will refer to different populations of cables based on whether the cable originates from a region of the nurse cell cortex adjacent to the follicle cells, the oocyte, or another nurse cell (Fig 1B,C). Some nurse cells have all cable types, and others only have nurse cables and follicle cables, depending on the location of a given nurse cell within the cluster (Fig 1B). We found that while nurse cables and oocyte cables are indistinguishable, follicle cables have distinct temporal growth dynamics. We show that this asymmetry in cable growth dynamics correlates with the distribution of the actin elongation factor Enabled (Ena); Ena localizes to the nurse cell cortex, but is excluded from the part of the cortex where follicle cables grow. In contrast, the formin Diaphanous (Dia) localizes to the nurse cell cortex at all interfaces. This suggested that Ena and Dia might collaborate to assemble nurse cables, while follicle cables might depend solely on Dia. Consistent with that model, we demonstrate that reduction of Dia impacted the initiation and elongation of both nurse cables and follicle cables, and that Ena was only necessary for nurse cable initiation and elongation. Mislocalization of Ena to the nurse cell cortex adjacent to the follicle cells eliminated the asymmetry in cable growth dynamics. Consistent with our model, we found that when cable growth dynamics were uniform, nuclear relocation was perturbed and dumping was incomplete. Together our data suggest that the differential localization of Ena and Dia is a key regulatory mechanism governing the asymmetry in cable growth dynamics. Further, our results support the model that cable growth asymmetry is necessary for the proper repositioning of the nurse cell nuclei and the successful completion of oogenesis.

## RESULTS

### Nurse cables, oocyte cables, and follicle cables exhibit a peak of initiation early in stage 10B and have similar densities

Using phalloidin to label fixed tissue and/or three different probes to detect actin in live tissue (SiR-Actin, and genetically encoded fTractin-tdTomato or LifeAct-GFP), we identified all types of actin cables at stage 10B (Fig.1 D-I). Previous studies showed that actin cables are not present on the follicle cell side of nurse cells in fixed egg chambers (Huelsmann et al., 2013), and we observed that as well (Fig. 1I). This suggests that follicle cables are more sensitive to fixation than nurse or oocyte cables. Actin cable sensitivity to fixation has been observed in other systems (Vasicova et al., 2016). While we visualized cables at all nurse cell interfaces using all three live imaging probes, we chose SiR-Actin for all subsequent experiments because it did not appear to induce any obvious changes to the actin cable array. In contrast, fTractin-tdTomato labelled actin cables had clusters of fluorescence at the base of each cable (Fig 1G, arrowhead) that we did not observe with SiR-Actin, LifeAct-GFP, or fixed. LifeAct-GFP appeared to induce higher densities of follicle cables (Fig 1H; Spracklen et al., 2014a) compared to SiR-Actin or fTractin-tdTomato.

To understand the dynamics and composition of the actin cable array, we examined the actin cable initiation rate during early stage 10B and the density of cables in different parts of the array. We were unable to image egg chambers at the end of stage 10A, prior to any cable initiation (Fig. 1A), as these chambers fail to progress in ex vivo culture (Spracklen and Tootle, 2013). Consequently, we isolated stage 10B egg chambers for imaging in the presence of SiR-actin, and selected for analysis those that were at the earliest points of cable initiation at the start of imaging. Individual egg chambers were imaged for a maximum of 90 minutes. Within a single nurse cell, we observed differences in the timing of actin cable initiation that depended on the z-axis position of the cables. For example, deeper nurse cables (Fig. 2A’-C’, yellow arrow) appear to initiate before more superficial cables (Fig. 2A-C, magenta arrow) as they are longer at 0 min. To assess initiation rate (Fig 2D,D’), we measured the number of new cables (<3 μm long) per μm of membrane at each time point from the beginning of stage 10B. For all cable types, the largest number of new cables appeared at the beginning of stage 10B (Fig. 2D). While the number of initiating cables significantly decreased for nurse and oocyte cables through the first 90 minutes of stage 10B (Fig. 2D’), initiation did not stop completely for any cable type (Fig. 2D). At time 0, the rate of cable initiation was significantly higher for nurse cables compared to follicle cables, but by 90 min the rate of new cable emergence was similar for all three cable types (Fig. 2D). Based on that rate of initiation through the first 90 min of stage 10B, we predicted that the cable density would significantly increase over this time (Fig. 2E, dotted lines). Surprisingly, while there was a trend toward increased cable density (Fig. 2E, solid lines), the density only increased significantly for follicle cables (Fig. 2E’). The dramatic difference between predicted and observed cable density is likely because not all cables that initiate continue to elongate (Fig. 2F,F’ yellow arrowheads). While there were some significant differences in cable density between the three cable types over the imaging period, by 90 min the densities of the three cable types were similar (Fig. 2E). Taken together, we observed some minor differences between the cable types in the rate of initiation and density, but overall these characteristics were very similar.

**Figure 2:**
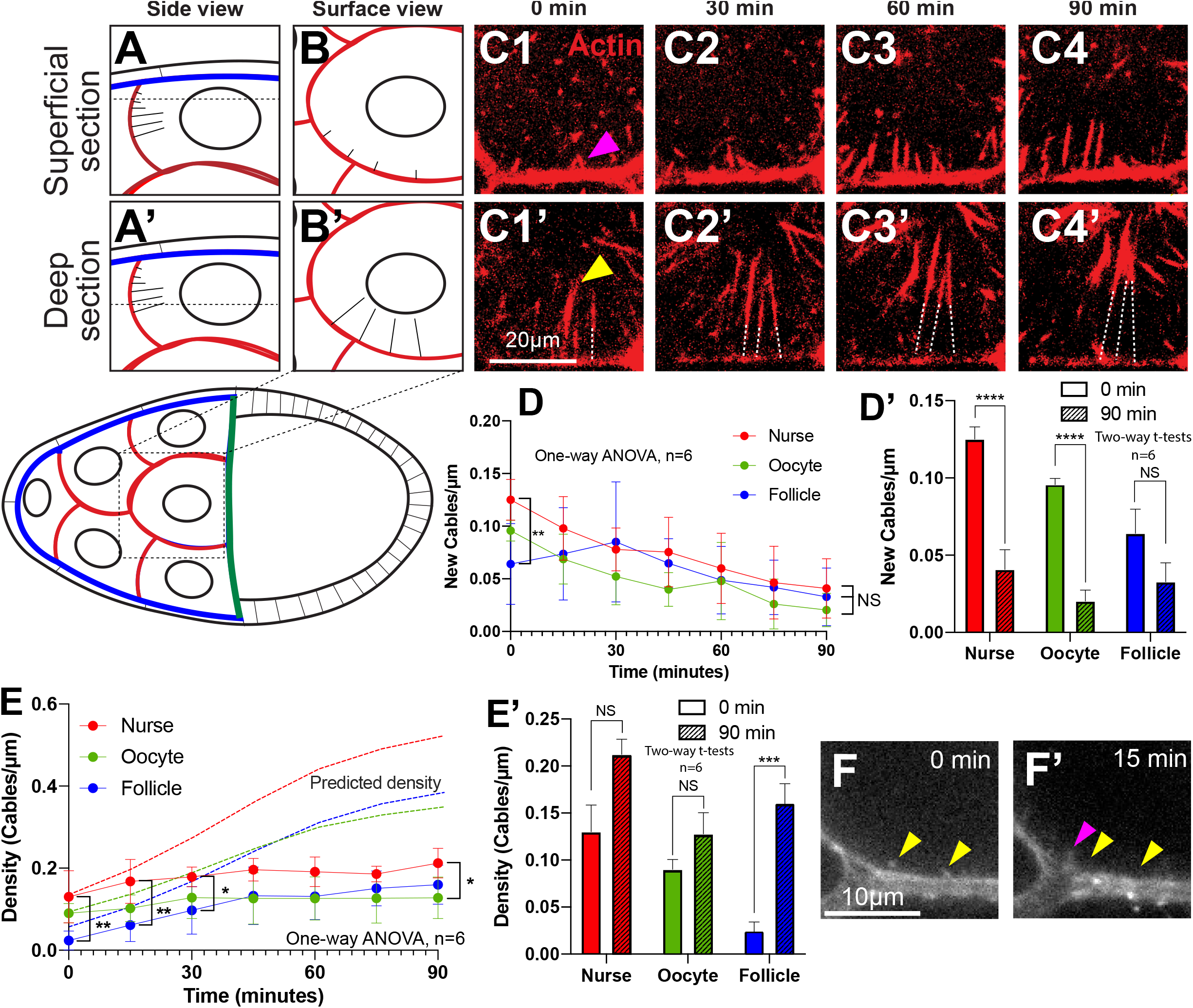
Cable types exhibit minimal differences in initiation rate and density. (A-C) Nurse cables growing from deeper planes initiate earlier than those from shallower planes. (A,B) Schematic of cables initiating from a superficial section of a nurse cell, closer to the follicular epithelium, shown in cross section (A), and in surface view (B). (C1-C4) Live imaging series of a surface view from a superficial section showing growth of SiR-Actin labelled cables (magenta). (A’,B’) Schematic of cables initiating from a deep section of a nurse cell, further from the follicular epithelium, shown in cross section (A’), and in surface view (B’). (C1’-C4’) Live imaging series of a surface view from a deep section showing growth of SiR-Actin labelled cables (yellow). These are significantly longer than those in the superficial section C1-C4. As the cables elongate they are often not in a single z-plane across their entire length, leading to apparent gaps near the cortex (dashed lines). (D) Actin cable initiation rate, measured as number of new cables (<3μm long) per μm of membrane at each timepoint starting at the beginning of stage 10B. (D’) Comparisons of cable initiation rate at the beginning (0 mins) and at the end (90 mins) of the imaging period. (E) Actual (solid lines) and predicted (dashed lines) actin cable density over time. (E’) Comparisons of actual cable density at the beginning (0 mins) and at the end (90 mins) of the imaging period. (F) Some cables initiate but appear to disassemble (yellow arrowheads). The magenta arrowhead indicates a newly initiating cable. *p<0.03, **p<0.002, ****p<0.0001. One way ANOVA with Tukey’s post-hoc analysis (D, E).

### Follicle cables grow slower and contact the nucleus later than nurse and oocyte cables

Because cables grow from all nurse cell cortices, we predicted that the different populations of cables would exhibit different growth rates to enable the cables to reposition the nuclei away from the ring canals during dumping. To test this, we measured the growth rate of each population of actin cables during the early stage of cable growth. Nurse and oocyte cables had a virtually identical growth rate of 0.11 μm/min (Fig. 3A,B). We observed the same growth rate for cables labelled with fTractin-tdTomato (data not shown). This was significantly faster than follicle cables, which grew at 0.07 μm/min (Fig. 3A,B). Stage 10B lasts for approximately 4 hrs, during which time the cables emerge and make contact with the nucleus. Therefore, we sampled the entire growth period in 90 min increments and integrated the data (see Methods) for the nurse cables (representative of the faster growing cables) and the follicle cables to determine if the growth rate changed over time (Fig. 3C). Both nurse cables and follicle cables maintained a constant growth rate throughout stage 10B (Fig. 3C). The nurse cell nuclei initiated movement between approx. 135 and 175 min. with dumping initiating between 210-240 min (Fig. 3C, shaded bars). Thus, both cable types grew at a constant rate from initiation through dumping.

**Figure 3:**
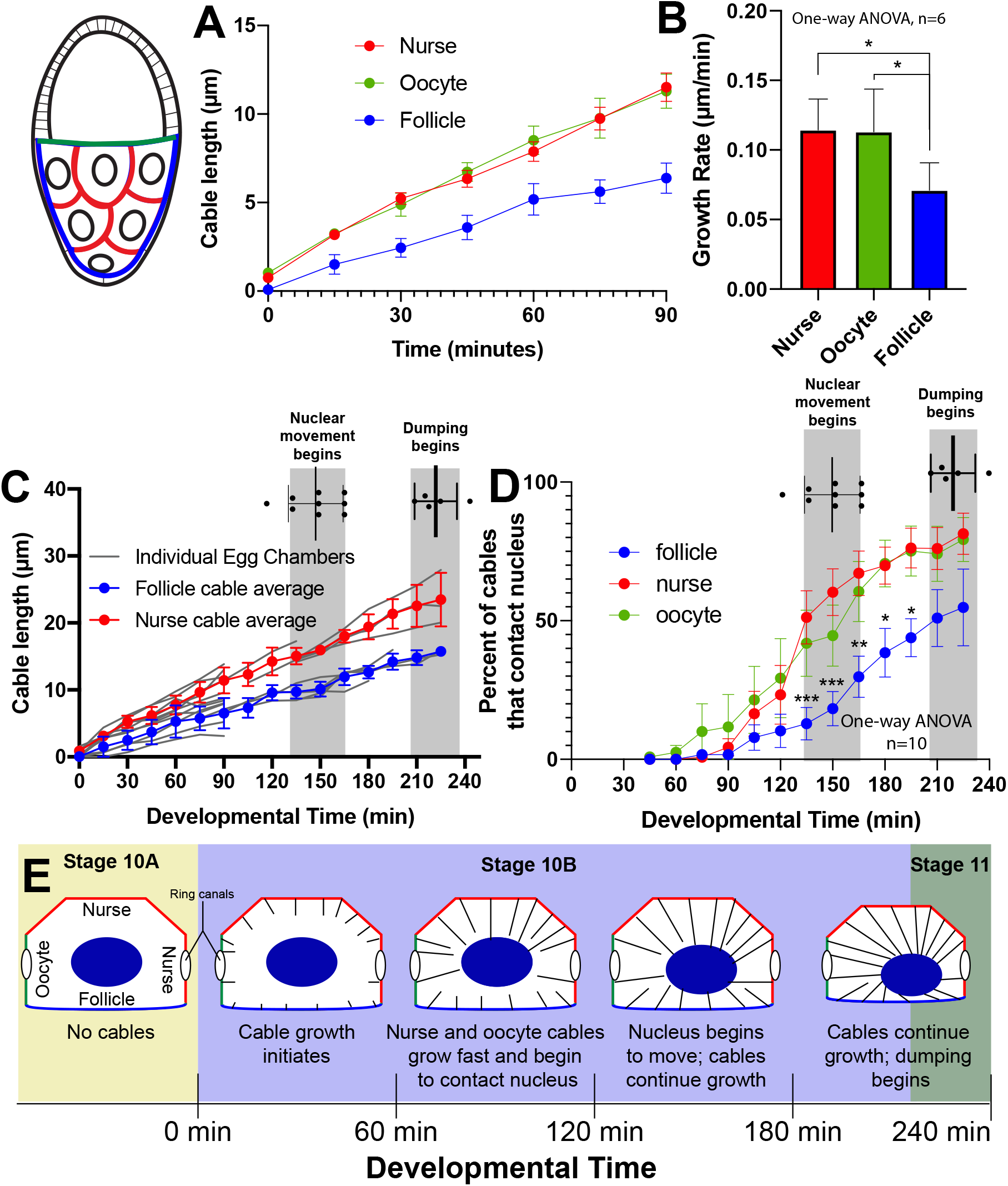
Cable types exhibit significant differences in growth rate and nuclear contact. (A-B) Follicle cable growth rate is significantly slower than that of nurse or oocyte cables. (A) Nurse, oocyte, and follicle cable length over the first 90 minutes of stage 10B. (B) Actin cable growth rate comparisons. (C) Nurse cable and follicle cable growth over the 4 hrs of stage 10B, showing consistent growth rates throughout. (D) Percent of cables that contact nuclei over time, showing that a significantly higher percentage of oocyte and nurse cables are in contact with the nucleus when the nucleus begins to move (first gray bar) compared to follicle cables. (E) Schematic showing a synopsis of the timing of actin cable growth through stage 10B illustrating the differences in growth rate and nuclear contact. The trajectory of nuclear movement is predicted. *p<0.03, **p<0.002, ***p<0.0002, Tukey’s post-hoc analysis.

Cables began to contact the nucleus 75-105 min. after the start of stage 10B with a steady increase in the percentage of cables making nuclear contact between 120 min and approximately 240 min (Fig. 3D). As nuclear movement began, the percent of nurse cables and oocyte cables contacting the nucleus was significantly higher than that of follicle cables (Fig. 3D). This suggests that the movement of the nuclei is primarily due to the activity of the nurse and oocyte cables. Taken together, our data demonstrate that cable growth rates and the dynamics of nuclear contact differ between different populations of actin cables. Nurse cables and oocyte cables, those cables that grow from the part of the cortex where the ring canals reside, grow faster and contact the nurse cell nuclei in greater numbers as the nuclei are moving than the slower growing follicle cables (Fig. 3E). These results support the model that spatial asymmetry in cable growth and nuclear contact repositions the nuclei away from the ring canals prior to dumping.

### Dia and Ena differentially localize to the nurse cell cortex

Our current model of actin cable growth is based largely on the EM study of Tilney and colleagues, and suggests that actin filament assembly factors present at the nurse cell cortex produce 2-4 μm actin filaments that are bundled into cable units that are then bundled into the growing cable (Guild et al., 1997). The spatial differences in cable growth rates that we observed suggest that different actin assembly factors may be responsible for nucleating and elongating the two different types of cables, the faster growing nurse and oocyte cables, and the slower growing follicle cables. The single Drosophila Ena/VASP family filament elongating protein, Enabled (Ena), was previously shown to be required for actin cable growth in nurse cells (Gates et al., 2007), and interacts with the formin Diaphanous (Dia) both in vitro and in vivo (Bilancia et al., 2014). Both Dia and Ena can independently promote the formation of filopodia, a structurally similar type of actin cable, but the dynamics and morphologies of those filopodia differ (Barzik et al., 2014; Bilancia et al., 2014; Homem and Peifer, 2008; Homem and Peifer, 2009; Nowotarski et al., 2014).

At stage 10B, we that found both Dia (Fig. 4A1-C1) and Ena (Fig. 4A2-C2) were enriched at the nurse cell cortex adjacent to other nurse cells (4B,D,E), and adjacent to the oocyte (Fig. 4F,G), i.e., at the sites of the fast-growing cables. In contrast, Dia (Figs. 4C1,D), but not Ena (Fig. 4C2,E), localized to the nurse cell cortex that is adjacent to the follicle cells where the slower growing follicle cables are found. These results suggest that Dia may assemble filaments independent of Ena to produce the slower growing follicle cables, while the faster growing cables may be the result of the combined activity of Dia and Ena. To examine this possibility further, we asked whether and how Dia and Ena colocalize at the nurse cell cortex. It was previously observed that the barbed, or growing end, of the cables are often found in membrane evaginations or “pits” that protrude into the neighboring nurse cell (Fig. 4H-K; Gates et al., 2009) The function of these cable pits is not known. Ena is prominently localized at the base of these pits (Fig. 4H1,4,I1,4, J; Gates et al., 2009), similar to its localization in the filopodial tip complex (Leijnse et al., 2015). Dia is largely uniform at the nurse cell cortex (Fig. 4H2,I2; yellow arrowhead), but also localizes to the tip of the cable pits with Ena, and along the shaft without Ena (Fig. 4H2,4, I4,4, J magenta arrowhead). In some filopodia, Ena and Dia exhibit a similar spatial relationship (Bilancia et al., 2014). On the follicle cell side of the nurse cell cortex, we observed Dia at the cortex, but did not find any evidence of cable pits (Fig 4L, magenta arrowhead). In summary, we find that Dia alone localizes to the part of the nurse cell cortex that grows the follicle cables, while Dia and Ena together localize in pits at the base of nurse cables (Fig. 4J,K). This suggests that the differential localization of Dia and Ena may be responsible for the differences in cable growth rates we observed.

**Figure 4:**
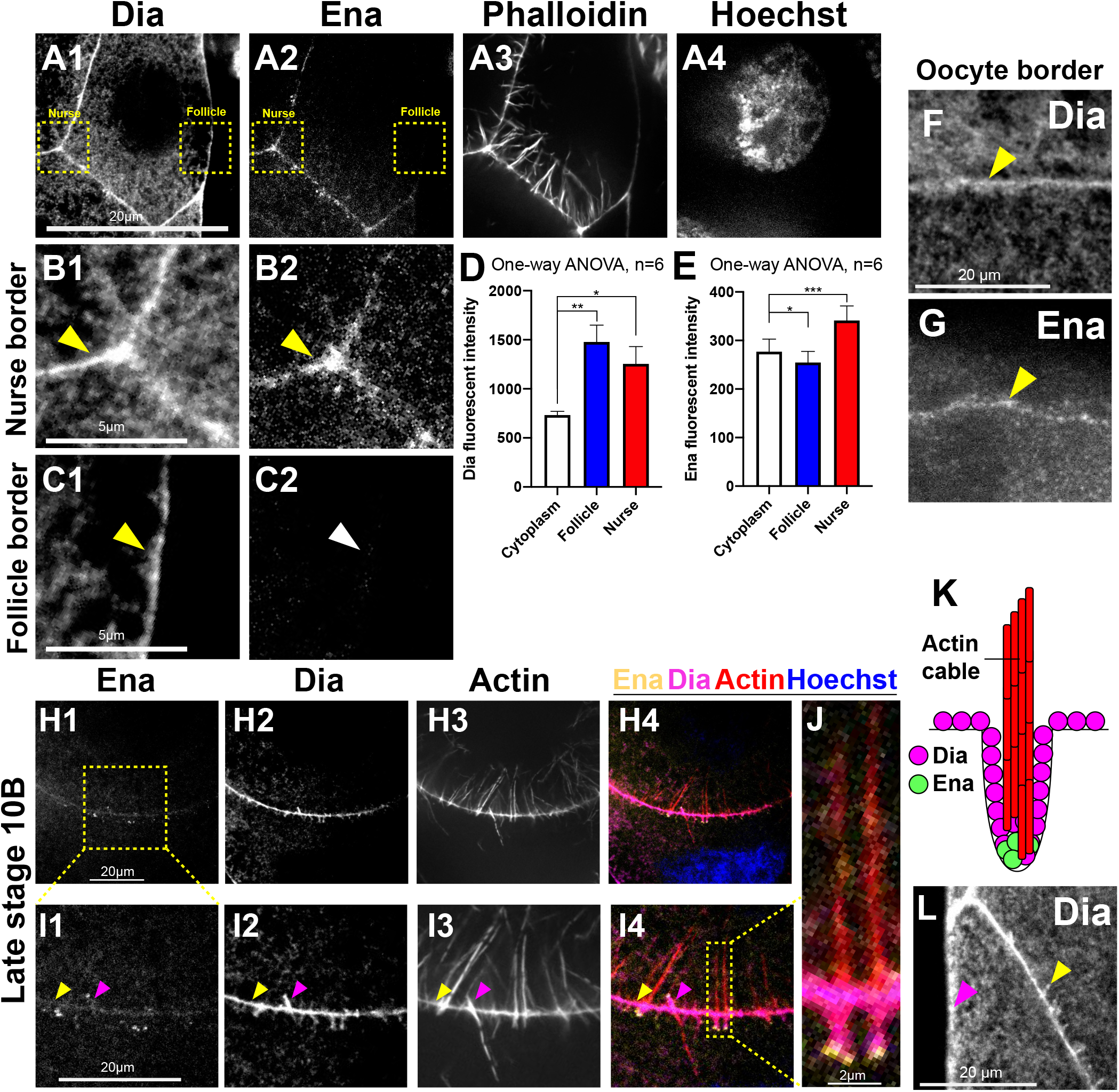
Ena and Dia have both overlapping and distinct localization patterns at the nurse cell cortex. (A-E) Dia is enriched in all regions of the nurse cell cortex, while Ena is excluded from the cortex bordering the follicular epithelium. (A-C) Images of fixed egg chambers labeled with anti-Dia, anti-Ena, phalloidin, and Hoechst. Boxed areas in A1-A2 are magnified in B1-C2 as indicated. Dia is enriched at the nurse cell cortex at both nurse cell-nurse cell borders (A1,B1, arrowhead), and at nurse cell-follicle cell borders (A1,C1 arrowhead). Ena is enriched at the nurse cell cortex at nurse cell-nurse cell borders (A2,B2, arrowhead), but not at nurse cell-follicle cell borders (A2,C2 arrowhead). (D) Quantification of Dia fluorescent intensity, showing Dia enrichment at both parts of the cortex. (E) Quantification of Ena fluorescent intensity, showing Ena enrichment only at nurse cell borders. Dia (F) and Ena (G) localize to the nurse cell-oocyte border. (H1-4) Ena and Dia localize to the actin cable pits. (I1-4) Higher magnification of indicated region in images H1-4 showing that Dia localizes along the pit shaft and base, while Ena localizes only to the pit base (magenta arrowhead). Dia is also generally enriched at the cortex (yellow arrowhead) (J) Higher magnification of indicated region in I4 showing actin cable pits. (K) Schematic of Dia, Ena, and actin localization in pits based on the image in J. (L) Dia localization to pits at nurse cell-nurse cell borders (yellow arrowhead), but not at nurse cell-follicle cell borders (magenta arrowhead). *p<0.03, **p<0.002, ***p<0.0002, Tukey’s post-hoc analysis.

### Reduction of Dia decreases nurse cable and follicle cable density, and reduces or halts cable growth

Our localization data suggested that Dia might be necessary for the growth of all nurse cell cable types. To test this, we reduced Dia levels in the nurse cells by driving the expression of two different *UAS-dia-dsRNA* transgenes using the *MDD-GAL4* (*MDD>*) maternal germline driver. Anti-Dia fluorescence intensity was significantly reduced at the nurse cell cortex at both nurse cell and follicle cell borders in tissue expressing either *dia-dsRNA* (Fig. S1A-E). Stronger Dia reduction using the *MTD-GAL4* (*MTD>*) driver blocked oogenesis prior to stage 10B. Consistent with our hypothesis, reduction of Dia resulted in a significant loss of both nurse and follicle cables (Fig. 5A-D,H). The remaining nurse cables often appeared in clusters (Fig. 5B,C, arrowheads), and exhibited growth defects (Fig. 5E). With both dsRNAs, approximately half of egg chambers had cables that exhibited a wild type growth rate, while the other half had slower growing or completely stalled cables (Fig. 5E-G). These types of nurse cable growth defects were very rarely observed in the *MDD>* control (Fig. 5E). Interestingly, all of the follicle cables remaining in the *dia-dsRNA* knockdown egg chambers had a significantly reduced growth rate compared to the *MDD>* control (Fig. 5H-J). This suggests that the follicle cables are more sensitive to Dia level, consistent with the absence of Ena on the follicle side of the cortex.

**Figure 5:**
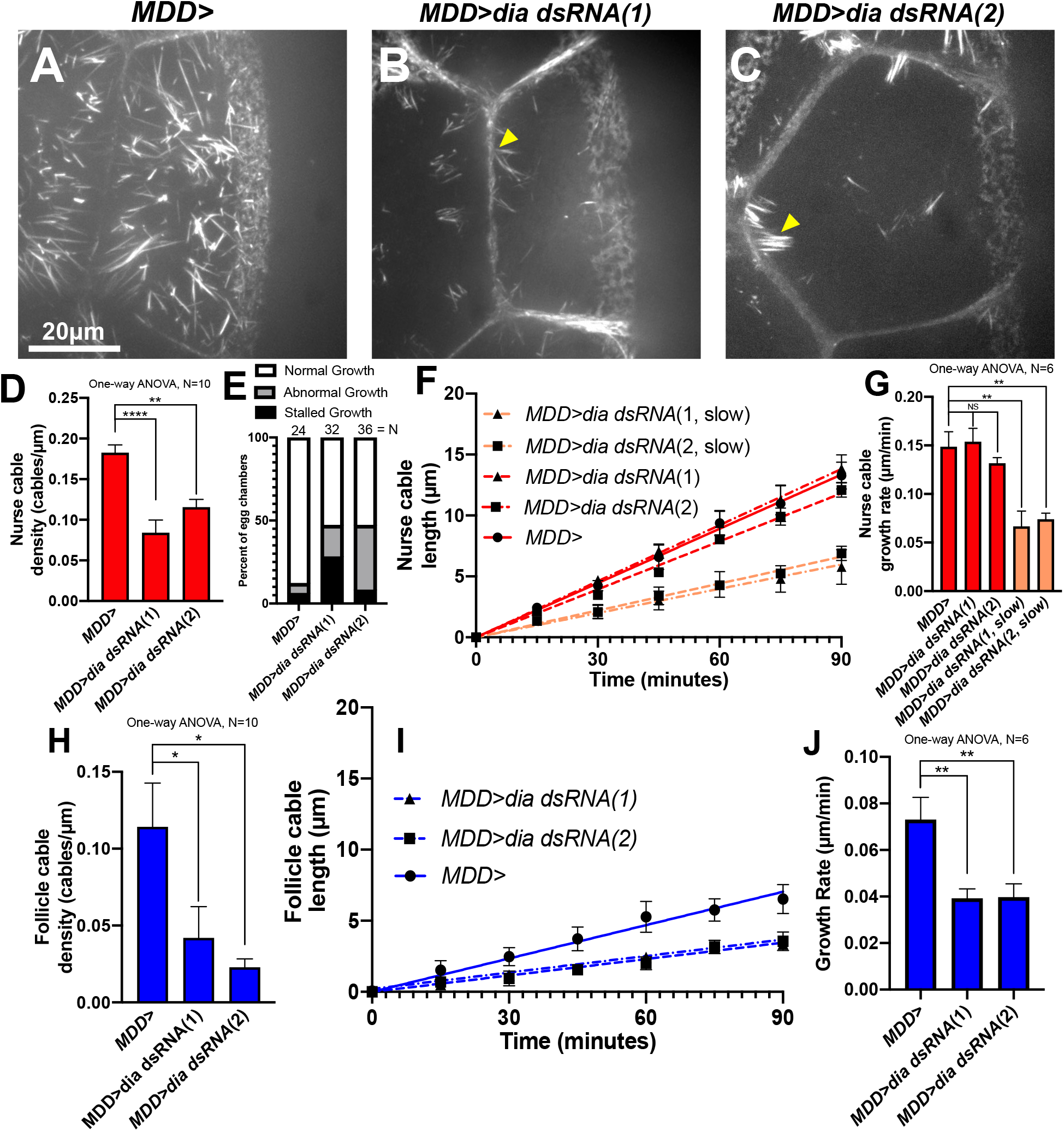
Dia is required for normal nurse cable and follicle cable initiation and growth rate. (A-C) Representative live images of actin cables in control (A), and two partial *dia-dsRNA* knockdown egg chambers (B,C), showing decreased cable density and clustered cables in the *dia* knockdown (arrowheads). (D) Quantification of nurse cable density, showing that both *dia* knockdowns have significantly lower cable density than controls. (E) Categorization of egg chamber cable growth phenotypes in genotypes as indicated. Reduction of Dia results in a higher proportion of egg chambers with abnormal or stalled cable growth. (F,G) Nurse cable length over time in control and *dia-dsRNA* egg chambers. A subset of nurse cables in *dia-dsRNA* egg chambers grow significantly slower than in the control. (H) The follicle cable density is significantly lower in *dia-dsRNA* egg chambers. (I,J) Follicle cable length over time in control and *dia-dsRNA* egg chambers (I). All follicle cables in *dia-dsRNA* egg chambers grow significantly slower than in the control (I,J).*p<0.03, **p<0.002, ***p<0.0002, ****p<0.0001, Tukey’s post-hoc analysis.

Although these phenotypes are consistent with the hypothesis that Dia is important for the growth of all nurse cell cables, the partial knockdown prevented us from determining whether formins are absolutely required for cable growth. To address this question, we added a formin inhibitor, SMIFH2 (Rizvi et al., 2009), to the medium at the start of imaging of stage 10B egg chambers. To determine how the inhibitor affected the growth rate, we assessed cable length over time (Fig 6A,C,H), and calculated the “instantaneous growth rate” over the 90 min imaging period by calculating the derivative of the nurse cable length and averaging the derivatives before and after each timepoint (Fig. 6B,D,I). This allowed us to compare the growth rates over time. By 30-45 min after the addition of 200 μM inhibitor, the cable growth rate was significantly reduced and after 45 minutes, the growth rate was no longer significantly different from 0μm/min (Fig. 6A,B). At 100 μM SMIFH2, nurse cable growth rate did not slow significantly, but cable growth stopped completely after 75 minutes (Fig 6A,B). Furthermore, when we simultaneously treated with 100 μM SMIFH2 and expressed *dia-dsRNA*, nurse cell cable growth decreased to zero immediately after treatment (Fig. 6C,D). These results strongly suggest that formins are necessary for nurse cable growth. In addition, we found that in the presence of 100 μM inhibitor the actin cables were contorted as they grew (Fig. 6E-G). In contrast to the dose dependent effects of SMIFH2 on nurse cable growth, the formin inhibitor had a faster and stronger effect on follicle cables (Fig. 6H,I). At both 100 μM and 200 μM, follicle cable growth stopped completely by approximately 15-30 min after the addition of inhibitor. Thus, formins are also necessary for follicle cable growth.

**Figure 6:**
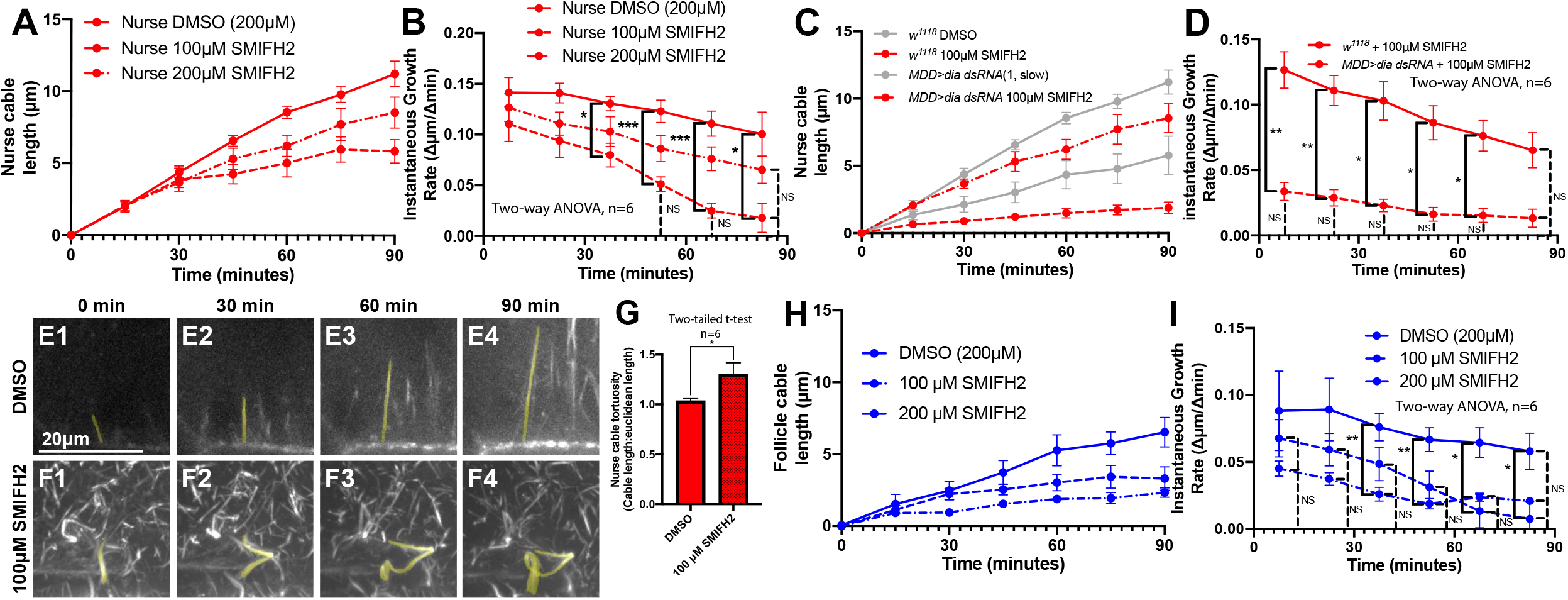
Formins are required for nurse cable and follicle cable growth. (A,B) Assessing the effect of the formin inhibitor SMIFH2 on nurse cable growth. (A) Nurse cable length over time after treatment with 100μM SMIFH2, 200μM SMIFH2, or DMSO. (B) Instantaneous growth rate calculated as the derivative of the curves in A. With the 200 μM treatment, the growth rate was not significantly different from 0μm/min at the last three time points (NS). (C,D) Assessing the effect of simultaneous *dia-dsRNA* knockdown and the formin inhibitor SMIFH2 on nurse cable growth. (C) Nurse cable length over time for *w^1118^* and *MDD>dia dsRNA* (1) egg chambers with and without 100μM SMIFH2. (Gray lines=data from Fig. 5F for comparison) (D) Instantaneous growth rate, calculated as derivative of curves in C. Treatment of *dia*-*dsRNA* knockdown egg chambers with the formin inhibitor blocked nurse cable growth. (E1-4) Linear nurse cable growth in a DMSO treated egg chamber. (F1-4) Bent nurse cable growth in wild type egg chamber treated with 100μM SMIFH2. (G) Quantification of nurse cable tortuosity from the conditions in E and F. (H,I) Assessing the effect of the formin inhibitor on follicle cable growth. (H) Follicle cable length over time after treatment with 100μM SMIFH2, 200μM SMIFH2, or DMSO. (I) Instantaneous growth rate of follicle cables calculated as derivative of the curves in H. At both inhibitor concentrations, follicle cable growth was blocked immediately. *p<0.03, **p<0.002, ***p<0.0002, Dunnett’s multiple comparisons post-hoc analysis (B,D,I).

Taken together, reduction of Dia activity by dsRNA knockdown and/or SMIFH2 inhibition dramatically reduced the number of nurse cables and follicle cables, and reduced or completely halted their growth. In all cases, the follicle cables were more sensitive to formin reduction/inhibition than nurse cables. This suggests that the slower growing follicle cables emanating from a Dia-positive, Ena-negative cortex are more highly dependent on Dia than the faster growing nurse cables emanating from a Dia-positive, Ena-positive cortex.

### Reduction of Ena decreases nurse cable density and growth rate, but does not disrupt follicle cable density or growth rate

Because Ena does not localize to the nurse cell cortex bordering the follicle cells, we predicted that Ena would play an important role in the growth of all cables except the follicle cables. To test this, we reduced Ena in the nurse cells by driving *ena-dsRNA* with a strong maternal germline driver (*MTD>*). This resulted in a significant decrease, but not complete loss, of Ena protein (Fig S1F-H). Consistent with our hypothesis, reduction of Ena significantly decreased nurse cable density (Fig 7A-C), with no effect on follicle cable density (Fig 7D-F). All of the remaining nurse cables in the nurse cells with reduced Ena exhibited a significant reduction in growth rate (Fig 7G,H), but the growth rate of the follicle cables was indistinguishable from the wild type control (Fig 7G,H). These results indicate that nurse cable initiation and growth are Ena dependent, but follicle cable initiation and growth are Ena independent.

**Figure 7:**
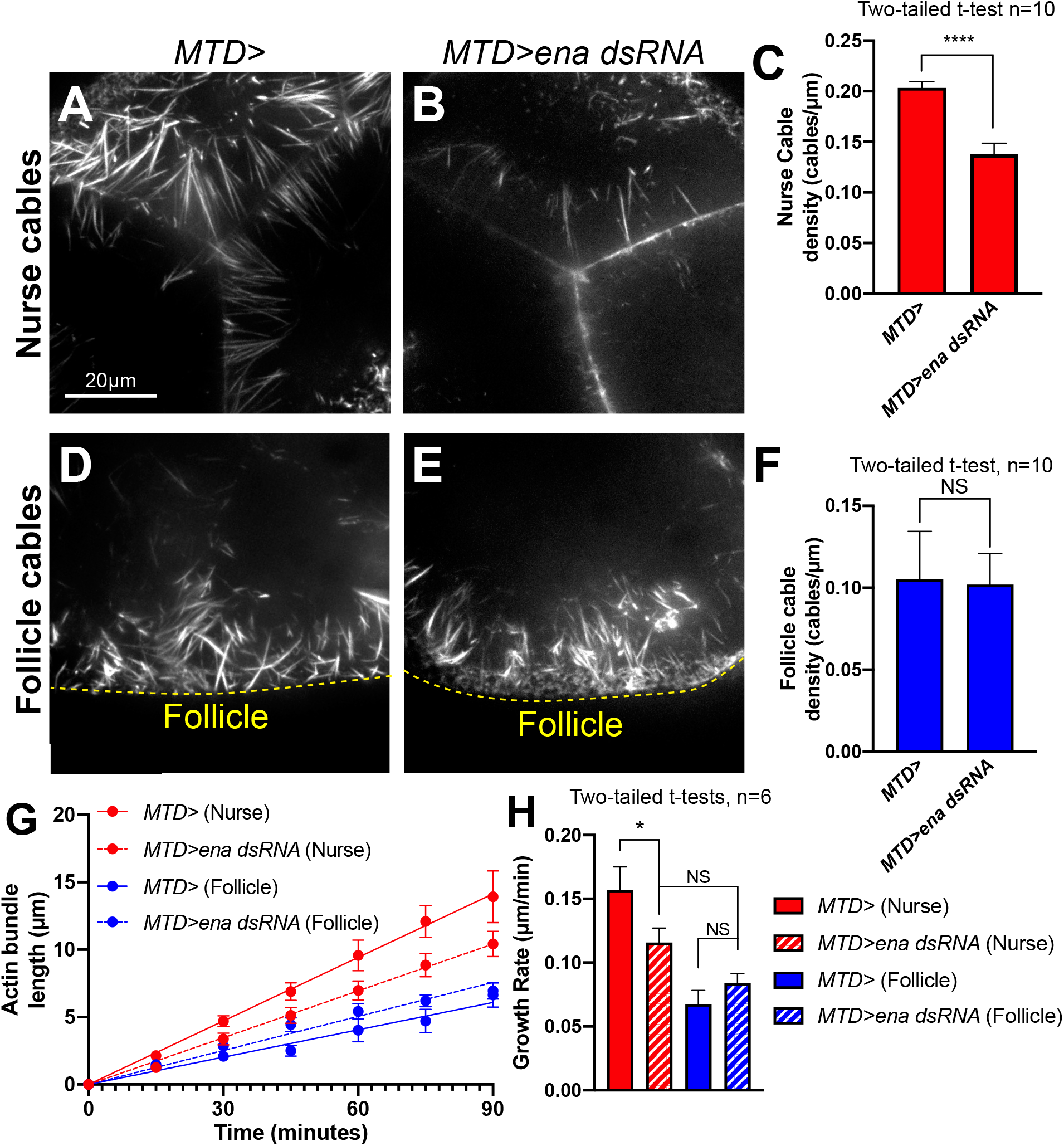
Ena is required for normal nurse cable initiation and growth, but is dispensable for follicle cables. Live images of nurse cables in control (A) and *ena* knockdown (B) egg chambers. (C) Nurse cable density is significantly lower in *ena* knockdown egg chambers compare to control. Live images of follicle cables in control (D) and *ena* knockdown (E) egg chambers. (F) Follicle cable density is not significantly different between control and *ena* knockdown. (G) Nurse and follicle cable length over time in control and *ena* knockdown egg chambers. (H) Nurse cables grow significantly slower when Ena is reduced, while follicle cable growth is unaffected. *p<0.03, ****p<0.0001.

### Mislocalization of Ena to the nurse cell cortex adjacent to the follicle cells creates uniform cable growth dynamics and prevents nuclear movement toward the follicle cells

Our model predicts that the spatial asymmetry in cable growth rate plays an important role in nurse cell nuclei relocation before dumping, and our data support the idea that the selective cortical localization of Ena is an important factor producing that asymmetry. We therefore asked whether mislocalization of Ena to the nurse cell cortex adjacent to the follicle cells would increase the rate of follicle cable growth and create uniform cable growth dynamics. To accomplish this, we overexpressed Ena by increasing the *ena* copy number using a stock containing 3 copies of *ena-mCherry* driven by the Ubiquitin promoter (3xEna-mCh). This overexpression resulted in Ena’s mislocalization to the nurse cell cortex adjacent to the follicle cells (Fig. 8A-B, arrowhead and insets). Consistent with our hypothesis, the follicle cable growth rate in 3xEna-mCh nurse cells significantly accelerated to match the rate of nurse cables in wild type or 3xEna-mCh nurse cells (Fig 8F,G). The uniform cable growth rate was also manifest in the overall cable length. At the end of stage 10B, follicle cables and nurse cables in 3xEna-mCh nurse cells were of similar length (Fig. 8D, E arrowheads), whereas nurse cables (Fig. 8C, magenta arrow) tend to be much longer than follicle cables (Fig. 8C, yellow arrow) in wild type nurse cells. Ena overexpression slightly, but not significantly, depressed the growth rate of wild type nurse cables (Fig. 8F,G). This may suggest that the mCherry-tagged Ena has partially reduced function compared to endogenous Ena. Regardless, the net result of Ena mislocalization was that the growth rate of the nurse and follicle cables matched the rate of wild type nurse cell cables, abolishing the normal asymmetry in growth rate (Fig. 8G).

**Figure 8:**
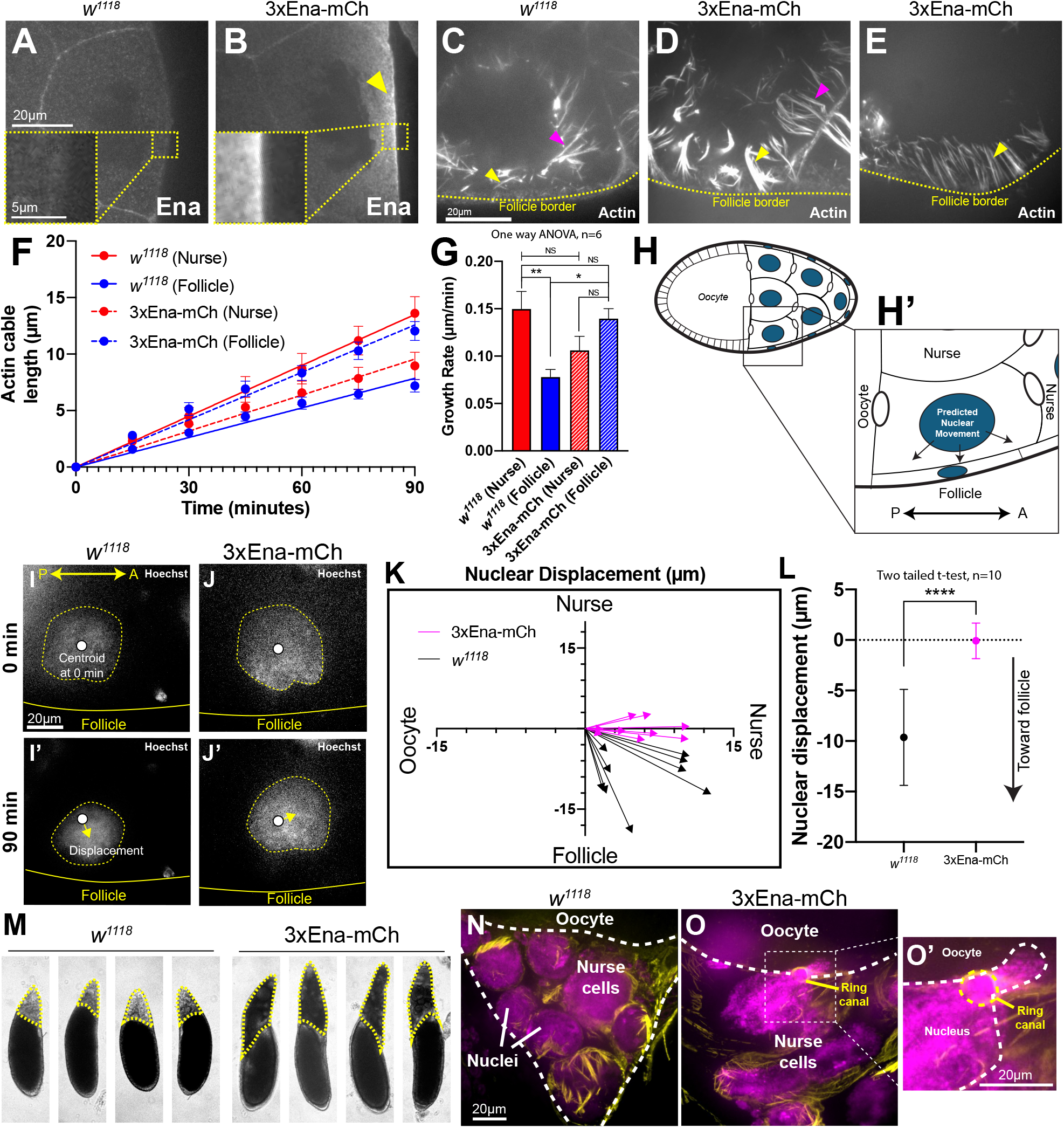
Mislocalization of Ena results in uniform cable growth dynamics, and defects in nuclear relocation and dumping. (A,B) Expression of 3xEna-mCherry (mCh) results in mislocalization of Ena to the nurse cell cortex at the follicle cell border (arrowhead). (C, D, E) 5μm z-projection of live, SiR-actin labelled egg chambers, showing nurse cables (magenta arrow) and follicle cables (yellow arrow) in wildtype (C) and 3xEna-mCherry (D,E) egg chambers. (F,G) Assessing the effect of Ena mislocalization on actin cable growth. (F) Nurse and follicle cable length over time in wildtype and 3xEna-mCh egg chambers. (G) Nurse and follicle cable growth rate in wildtype and 3xEna-mCh egg chambers. Follicle cable growth rate is significantly increased in 3xEna-mCh egg chambers. (H,H’) Schematics of the position of the nurse cell that we selected to measure nuclear movement, depicting the predicted wild type nuclear trajectory. (I) Live wildtype egg chamber stained with Hoechst, showing that over 90 min the nucleus moves toward the follicular epithelium (I’). (J) Live 3xEna-mCh egg chamber stained with Hoechst, showing that over 90 min the nucleus moves toward the neighboring anterior nurse cell (J’). (K) Nuclear displacement distance in wildtype (black) and 3xEna-mCh (magenta) egg chambers over 90 min (n=10). (L) There is no significant nuclear displacement toward the follicular epithelium in 3xEna-mCh egg chambers (magenta). (M) Representative brightfield images showing nurse cells (yellow dashed lines) that do not complete dumping in 3xEna-mCh egg chambers compared to wild type. (N,O) Representative immunofluorescence images stained with Hoechst (magenta) and phalloidin (yellow) of stage 12 wildtype (N) and 3xEna-mCh egg chambers showing an example of nuclear extrusion (O,O’). **p<0.002, ****p<0.0001, Tukey’s post-hoc analysis (G).

We next asked whether the uniform cable growth rate would prevent the normal displacement of nurse cell nuclei prior to dumping. The position of a nurse cell in the 15 cell syncytium determines its neighbor relationships and affects the position of the ring canals in that cell. To control for this variability, we characterized nuclear movement in nurse cells bordering the oocyte and the squamous follicular epithelium (Fig. 8H,H’). Based on this geometric arrangement and our analysis of cable growth dynamics, we predicted that as these nuclei would move toward the follicle cells (Fig. 8H’). While the precise trajectory and final location varied, all nuclei in wild type egg chambers moved toward the nurse cell cortex at the follicle cell border, or toward the boundary between the anterior nurse cell and the adjacent follicle cells (Fig. 8I,K,L black). When we equalized actin cable growth rate by mislocalizing Ena, we found that the nuclei were no longer displaced toward that region of the nurse cell. Instead, nuclei in cells overexpressing Ena-mCherry moved exclusively toward the anterior nurse cell and consequently toward the ring canal at that boundary (Fig. 8J-L, magenta). This demonstrates that asymmetric cable growth in wild type nurse cells is required for the proper repositioning of nuclei prior to dumping.

We predicted that the defect in nuclear displacement we observed in this specific population of nurse cells would also be true in the other nurse cells impeding cytoplasmic dumping. To test this, we assessed dumping by examining stage 12 egg chambers from control and 3xEna-mCh ovaries. 90% (n=40) of wild type egg chambers completed dumping, resulting in a small remnant of the nurse cell cluster (Fig 8M, dashed lines). In contrast, only 13% (n=40) of 3xEna-mCh egg chambers completed dumping, and in the remaining egg chambers we observed a large nurse cell remnant (Fig. 8M, dashed lines). We predicted that nuclear blockage of ring canals resulting from defects in nuclear displacement would contribute to the dumping defect. Consistent with this, we found examples of nuclear extrusion through the ring canals in 3xEna-mCh egg chambers (Fig. 8O,O’) reminiscent of what was observed in Singed/Fascin and Quail/Villin mutants (Cant et al., 1994; Mahajan-Miklos and Cooley, 1994). We never observed nuclear extrusion in *w^1118^* egg chambers (Fig. 8N). The anomalous dumping we observed could be attributed exclusively to nuclear relocation defects, but we cannot rule out the possibility that defective nurse cell cortical contractility resulting from Ena overexpression and mislocalization may also contribute. Together our results support the model that asymmetric actin cable growth rate is critical for proper nuclear displacement in nurse cells to enable the successful completion of cytoplasmic dumping and oogenesis.

## Discussion

Nurse cell nuclei are moved away from the ring canals in a late stage of oogenesis to facilitate the rapid transport of nurse cell cytoplasm into the oocyte. Early work in fixed tissue suggested that the cable array was a static basket, holding the nuclei in place while the nurse cell cytoplasm flowed past through the ring canals (Cant et al., 1994; Mahajan-Miklos and Cooley, 1994). Live imaging revealed that instead, the actin cable array actively contacts and pushes the nuclei, rolling and wrapping them as the cables elongate (Huelsmann et al., 2013). Here we show that the actin cable array is not uniform, and that the asymmetry in cable growth rate we discovered is necessary for the directional repositioning of nurse cell nuclei and the completion of dumping. Further, we show that both Dia and Ena are required to initiate and elongate the faster growing nurse cables, while Dia alone is necessary for the proper initiation and elongation of the slower growing follicle cables. Finally, our results suggest that selective localization Ena is a key factor that establishes the asymmetry in the dynamic properties of the actin cable array.

The use of an asymmetric actin cable array to push and relocate nuclei is novel amongst the diversity of actin-based mechanisms for nuclear movement. Actomyosin contraction (Nakazawa and Kengaku, 2020; Sakamoto et al., 2020) and actin flows (Zhu et al., 2018) move nuclei in migrating cultured cells and neurons. Active diffusion of actin coated vesicles driven by myosin Vb generates a pressure gradient and propulsive force to move the oocyte nucleus in the mouse (Almonacid et al 2015). Actin cables are utilized in other contexts, but in those cases nuclei are coupled to the actin cables used as tracks for nuclear transport (Folker et al., 2011; Luxton et al., 2010; Luxton et al., 2011). The retinal neuroepithelium in fish utilizes a different sort of formin-dependent actin network to push the nucleus apically, and cortex-anchored and bundled actin modulated by myosin may generate the pushing force (Yanakieva et al., 2019). Microtubule dependent pushing and pulling forces are also widely used to move nuclei (Adames and Cooper, 2000; Gönczy et al., 1999; Levy and Holzbaur, 2008; Tran et al., 2001; Zhao et al., 2012).

Our data suggest the actin cable array pushes nuclei directionally as the result of the asymmetry in cable growth rates: the faster growing nurse and oocyte cables make first contact with the nucleus and begin to push it toward the slower growing cables on the follicle cell side of the nurse cell cortex (Fig 3E). This is consistent with the overall trajectory of the nucleus in our sample nurse cell that was toward the follicular epithelium and away from the ring canals (Fig 8K). When we equalized cable growth rate through the mislocalization of Ena, the nuclei embarked on a distinct trajectory toward the ring canal connecting to the neighboring anterior nurse cell (Fig 8K). We were surprised that the nucleus moved at all-if the actin cable array was uniform in all features one might predict that the nucleus would be trapped in its original position by equivalent force exerted on all sides of the nucleus. Stochastic differences in the “uniform” cable array would be predicted to result in random movement of the nucleus, contrary to the directional movement we observed (Fig. 8K). This suggests that the cable array with apparently uniform growth rate may harbor a more subtle asymmetry that contributed to the directional, though misguided, nuclear movement.

The nurse cell actin cables share structural similarity with Listeria tails that contain crosslinked bundles of actin filaments (Jasnin et al., 2013), and with filopodia that typically contain a single bundle of 10-30 unidirectional linear actin filaments extending from a few to 100 μm (Leijnse et al., 2015). Cables can be simply viewed as inverted filopodia where the actin bundles extend into the cytoplasm rather than away from the cell body. Despite this difference in orientation, cables and filopodia share a common set of assembly factors and bundlers. Both utilize formins and Ena/VASP proteins to assemble filaments bundled by Fascin and Villin (Gates et al., 2009; Huelsmann and Brown, 2014; Leijnse et al., 2015; Figs. 5–7). In filopodia, Ena localizes to a well described “tip complex” at the distal end of the filopodium that elongates away from the cell body. Ena similarly localizes to the base of nurse cables and oocyte cables, the site of filament assembly (Guild et al., 1997), that project outward 1-2 μm in cable pits that protrude into the neighboring nurse cell (Gates et al., 2009; Guild et al., 1997; Fig 4J,K). We have shown that Dia also localizes to these pits, where it can be found along the shaft of the pit and colocalized with Ena in the tip (Fig. 4J,K). Similar to what we observe in cable pits, endogenous Ena and Dia can sometimes be found associated with the same filopodium with Ena at the tip and Dia in the shaft (Bilancia et al., 2014).

The functional relationship between formins and Ena/VASP in filopodia is a complex one. Independently, they create filopodia and other projections with distinct morphological, dynamic, and functional properties (Barzik et al., 2014; Bilancia et al., 2014; Homem and Peifer, 2008; Homem and Peifer, 2009; Nowotarski et al., 2014). For example, Bilanica et al. (2014) showed that in Drosophila D16 cultured cells, Ena overexpressed alone promoted the formation of filopodia that exhibited dynamic changes in number, length and lifetime. In contrast, Dia overexpressed alone promoted the formation of long, stable filopodia. These in vivo activities may be explained in part by their different in vitro activities, where Dia is a faster and more processive elongator. Because Ena is slower and less processive, other actin modifiers may have the opportunity to alter the characteristics of the developing filopodia. When Dia and Ena were co-overexpressed, the cells produced fewer, shorter filopodia, suggesting that Ena inhibits Dia. Consistent with that, the Ena EVH1 domain binds Dia and inhibits its nucleation activity in vitro. Furthermore, in the very small number of filopodia where Dia and Ena colocalized in the tip complex, the majority retracted, folded back, or stalled.

Our data suggest that Dia and Ena interact to assemble nurse cables and oocyte cables, but the functional consequence of the interaction appears to be different from what has been observed in filopodia. First, we find that endogenous Dia consistently colocalizes with Ena in the cable “tip complex” (Fig 4H,I), suggesting that colocalization does not terminate cable growth as predicted from the work of Bilancia et al. (2014). Further, our functional data are most consistent with a collaboration between Dia and Ena in both cable initiation and elongation, as reduction/inhibition of either protein significantly reduced cable number and slowed or halted nurse cable growth (Figs 5–7). Interestingly, the nurse and oocyte cables are also faster growing than the follicle cables where Dia functions without Ena (Figs 5–7). The previous work described above would predict the opposite (Bilancia et al., 2014). This may suggest that the Dia-Ena collaboration accelerates filament assembly, or that there are other factors modulating the rate of cable growth of one or both cable types.

Our model suggests that the selective localization of Ena to the nurse cell cortex adjacent to other nurse cells and adjacent to the oocyte is a key factor in generating the asymmetric actin cable array. Adherens junctions between nurse cells, and between the nurse cells and oocyte, is one way in which this cortical region may differ from that adjacent to the follicle cells. The nature of physical interactions between the nurse cells and overlying squamous follicle cells is not clear. One potential regulator of Ena localization is prostaglandin signaling, which contributes to proper Ena localization during stage 10B (Spracklen et al., 2014b), and is required for nurse cell dumping (Tootle and Spradling, 2008). Another candidate, Lamellipodin (Lpn, Drosophila Pico), is important for the localization of Ena to lamellipodia in mammalian cells (Carmona et al., 2016; Cheng and Mullins, 2020; Hansen and Mullins, 2015; Michael et al., 2010), and also contributes to nurse cell dumping (Spracklen et al., 2019). Loss of Abelson (Abl), a tyrosine kinase targeting Ena, results in a dumping defect, premature cable assembly, and some abnormal Ena localization (Gates et al., 2009). Ena’s selective cortical localization in epithelia and other cell types is also though to influence protrusive activity and interactions with Dia (Bilancia et al., 2014; Gates et al., 2007; Homem and Peifer, 2009; Nowotarski et al., 2014). Thus, tight control of Ena’s cortical localization may be a common mechanism regulating actin filament assembly that impacts a wide variety of processes from cell migration to morphogenesis and nuclear positioning.

## Acknowledgements

Thank you to Haibing Teng and the Molecular Biosensor and Imaging Center (MBIC) at Carnegie Mellon University for providing us with some imaging resources. The anti-Dia antibody was a gift from S. Wasserman (UCSD). Many thanks to C. Ettensohn for valuable comments on the manuscript. This work was funded by a grant to B.M. from the National Institutes of Health (R01-GM120378). There are no competing interests.

## Methods

### Flies

The following stocks were used: *w^1118^, MDD-GAL4* (Bloomington #80361), *MTD-GAL4* (Bloomington #31777), *UAS-Ftractin-tdTomato* (Bloomington #58989), *UAS-LifeAct-GFP* (Bloomington #57326), *UAS-dia-dsRNA*(1) (HMS00308, Bloomington #33424), *UAS-dia-dsRNA*(2) (GL00408, Bloomington #35479), *UAS-ena-dsRNA* (Bloomington #35479), *Ubi-ena-mCherry* (Bloomington #58731). 24 hours before all experiments, flies were fed wet yeast paste to promote egg production.

### Imaging

For live imaging experiments, stage 10B were isolated as described (Spracklen and Tootle, 2013). Egg chambers were imaged in live imaging media consisting of Schneider’s media with 20% FBS, 5 μg/mL insulin, 2 mg/mL trehalose, 5 μM methoprene, 1 μg/mL 20-hydroxyecdysone, and 50 ng/mL adenosine deaminase in Concanavalin-A coated glass bottom petri dishes (Azer Scientific ES56291). At the start of imaging, egg chambers were labeled with the nuclear stain 1:5000 SYTO83 (Thermofisher, S11364) and/or 1:1000 SiR-actin (Spirochrome, CY-SC001, Stein am Rhein, Switzerland). For *w^1118^* actin cable characterization experiments (Figs. 2,3), we used an Andor Revolution XD spinning disk confocal microscope. All other images were acquired with a Prime95B CMOS camera (Photometrics) on a Zeiss Axiovert 200M microscope with a X-Light V2 spinning disc scan head (Crest Optics). Fluorophores were excited with a Celesta Light Engine (Lumencor). For live imaging, stacks (z-step 0.5μm) were taken every 15 minutes for 90 minutes. For formin inhibition experiments, SMIFH2 (Sigma, S4826) or DMSO was added directly to the imaging media at the beginning of the imaging period. Ovaries were fixed for 20 min. in 4% paraformaldehyde diluted in phosphate buffered saline (PBS) from an 8% stock (EMS #157-8) and blocked for 1 hr in PBS plus 0.2% Triton X-100 and 4% normal goat serum (NGS). Antibody/label incubations were in PBS plus 0.1% Triton X-100 and 1% NGS using 1:500 Alexa488-Phalloidin (ThermoFisher, A12379), 1:1000 Hoechst (Invitrogen, 33342), and the following antibodies: 1:1000 mouse anti-Ena (DSHB, 5G2) and 1:1000 rabbit anti-Dia (gift from Wasserman lab, UCSD).

### Quantification of actin cable properties and growth dynamics

To quantify cable initiation and density, where it is critical to detect cables in early stage 10B, verapamil (1:1000) was added to the media to increase the rate of SiR-actin labeling. For all experiments, n refers to the number of egg chambers. Where relevant, 5 cables per egg chamber were analyzed. For **initiation measurements**, the number of cables less than 3μm long was measured at each timepoint along 2-3 spans of membrane per egg chamber totaling a distance of 120-240μm for nurse borders, 70-100μm for oocyte borders, and 60-140μm for follicle borders. The predicted density was measured as the cumulative sum of all new cables on and before each timepoint. For **density measurements**, the total number of actin cables was measured over 5 spans of nurse cell borders totaling 150-240μm in length, 2-5 spans of oocyte borders totaling 60-220μm in length, and 2-3 spans of follicle borders totaling 70-170μm in length. For **cable growth rates**, the length of five actin cables in each of 6 egg chambers was tracked over the imaging period and linear regressions were taken to determine the growth rate of each cable. To measure **time of dumping or nuclear movement**, we measured the length of 10 nurse cables from egg chambers at the beginning of dumping (marked by decreasing nurse cell size) or from egg chambers at the beginning of nuclear movement; then we calculated the time of dumping or nuclear movement by dividing the average cable length of each egg chamber by the average nurse cell growth rate. To determine the percent of cables that contact the nucleus, we assessed nuclear contact of 10-20 randomly chosen cables from 2-3 nurse cells per egg chamber. We calculated Dia and Ena intensity in fixed egg chambers by measuring average intensity along 5 segments (~5μm long) of nurse border, follicle border, or cytoplasm in each of 6 egg chambers. For **tortuosity** measurements, we measured the actual length of 10 cables and divided by the Euclidean distance of each cable (linear distance between the barbed and pointed ends). When measuring growth rate for experiments where it is not critical to include the beginning of stage 10B, we subtracted the length of each cable by the length of the cable at t0; this allowed for uniform growth curves with the length of each curve starting at 0μm. Instantaneous growth rate was calculated using the derivative tool in Prism 8, and smoothing the resulting curve by averaging each value with its neighboring values.

### Quantification of nuclear displacement

All nuclear displacement measurements were done in nurse cells directly adjacent to the oocyte, and images were rotated or flipped so the oocyte was to the left of the nurse cell and the follicle was to the bottom. To take movement of the egg chamber into account, we determined cell-normalized nuclear location by subtracting the centroid coordinates of the nucleus by the centroid coordinates of the nurse cell at 0 minutes and 90 minutes. For graphical representation, we generated vectors from the centroid at 0 minutes to the centroid at 90 minutes and transformed each vector to the origin of the graph. The displacement toward the follicle is the displacement along the y-axis of each vector. We followed nuclei for 90 minutes from the start of imaging, selecting for analysis only egg chambers at the developmental stage where the nucleus was expected to move, which we identified as the point where all cables begin to contact the nucleus (Fig. 3D).

### Image analysis and statistics

We performed all measurements using ImageJ (version 2.0.0). All analysis and graphs were generated with Prism 8 (Graphpad), and figures were prepared using Adobe Illustrator.

